# Prediction of Four Novel SNPs V17M, R11H, A66T, and F57S in the *SBDS* Gene Possibly Associated with Formation of Shwachman-Diamond Syndrome, using an Insilico Approach

**DOI:** 10.1101/542654

**Authors:** Anfal Osama Mohamed Sati, Sara Ali Abdalla Ali, Rouaa Babikir Ahmed Abduallah, Manal Satti Awad Elsied, Mohamed A. Hassan

**Affiliations:** Department of Applied Bioinformatics - Africa City of Technology, Biotechnology Park, Khartoum- Sudan

**Keywords:** Shwacman Diamond Bodian, Shwachman Diamond Syndrome, SDS, SBDS gene, Single Nucleotide Polymorphisms, SNPs, mutations, Bioinformatics, insilico, bioinsilico, single nucleotide variations

## Abstract

Shwachman-Diamond syndrome SDS (MIM 260400) is an autosomal recessive disorder. Characterized by exocrine insufficiency of the pancreas, bone marrow hypoplasia resulting in cytopenias, especially neutropenia, variable degree of skeletal abnormalities, failure to thrive, and increasing risk of developing myelodysplasia or transformation to leukemia. SDS is mainly caused by Shwachman Bodian Diamond Syndrome gene (MIM ID 607444). SBDS gene is 14899 bp in length, located in chromosome seven in the eleventh region of the long arm, and is composed of five exons. It’s a ribosomal maturation factor which encodes a highly conservative protein that has widely unknown functions despite of its abundance in the nucleolus A total number of 53 SNPs of homo sapiens SBDS gene were obtained from the national center for biotechnology information (NCBI) analyzed using translational tools, 7 of which were deleterious according to SIFT server and were further analyzed using several software’s (Polyhen-2, SNPs&Go, I-Mutant 2.0, Mutpred2, structural analysis software’s and multiple sequence alignment software). Four SNPs (rs11557408 (V17M), rs11557409 (R11H), rs367842164 (A66T), and rs376960114 (F57S)) were predicted to be disease causing, localized at highly conservative regions of the SBDS protein and were not reported in any previous study. In addition, the study predicts new functions of the SBDS protein DNA related, and suggests explanations for Patients developing cytopenias and failure to thrive through genetic co-expression and physical interaction with RBF1 and EXOSC3, respectively.

## Introduction

Shwachman-Diamond syndrome (SDS; MIM 260400) is an autosomal recessive disorder. Characterized by exocrine insufficiency of the pancreas, bone marrow hypoplasia resulting in cytopenias, especially neutropenia, variable degree of skeletal abnormalities, failure to thrive, and increasing risk of developing myelodysplasia or transformation to leukemia. Other common features were found in younger patients. Such as, hepatomegaly, abnormal liver and biochemical tests findings.^1–13^ SDS is a rare disorder which counts for 1:77000 individuals^14^. The hematological abnormalities was a constituent in 88% of the cases. Neutropenia in 95%, anemia in 50%, and thrombocytopenia in 70% of cases. 50% of the cases presented with short stature and severe infections occurred in 85% of cases. In addition, some degree of mental retardation and learning disabilities was reported in some of the studied cases. ^10,15–18^

SDS is mainly caused by Shwachman Bodian Diamond Syndrome gene (*SBDS*, gene bank ID 51119). ^12,15,16,19,20^ *SBDS* gene is 14899 bp in length, located in chromosome seven in the eleventh region of the long arm, and is composed of five exons^21,22^. *SBDS* gene is a ribosomal maturation factor encodes a highly conservative protein which has widely unknown functions despite of its abundance in the nucleolus. However, in SBDS orthologs it’s known to play a vital role in ribosomal biogenesis. ^20,21,23–27^SBDS along with the 60S large ribosomal subunit migrate and precipitates with 28S ribosomal RNA (rRNA). However, loss of SBDS Protein has not been reported to be associated directly with an individual block in rRNA maturation. Yet, there’s a notable increase in immature 60S ribosomal subunits.^5,26^ Additionally, SBDS forms a protein complex with nucleophosmin, a protein involved in maintenance of genomic integrity, ribosomal biogenesis and generation of leukemia.^26,28,29^

Shwachman-Diamond syndrome is an interesting disease with a high probability of transitioning to leukaemia and myelodysplasia and a better understanding of the clinical aspects of this rare disorder would assist in providing further information about its pathology. Many previous studies had various approaches to understand the pathophysiology of the *SBDS* gene and the possible impact of its chromosomal aberrations in the formation of the disease.^3,20,21,23,26,30–32^ However, the key answer to these questions remains unknown. Therefore, the aim of this study, is to analyze the single nucleotide polymorphisms (SNPs) of the SBDS gene to unveil any possible relationships between single amino acid substitution mutations (missense mutations) and development of SDS.

In this paper, translation technology was used to analyze SNPs of the SBDS gene and assess both gene and protein functions and the impact of these mutations on gene expression. This study is different than other studies for being the first to use computational tools to analyze SBDS gene SNPs.

### Methodology

A total number of 51 SNPs of Homo sapiens SBDS gene were obtained from the National Center for Biotechnology Information (NCBI) on October 2018, of which twenty-four SNPs were in the 5’ and 3’ un-translated regions, and twenty-seven non-synonymous SNPs (nsSNPs) were in coding regions. The latter, was analyzed using several softwares with different approaches and working algorithms, this is to ensure accuracy of results as insilico tools work consistently, and thus, one tool compensates for other tool’s weaknesses.(Figure 1)^33^

**Figure (1):**
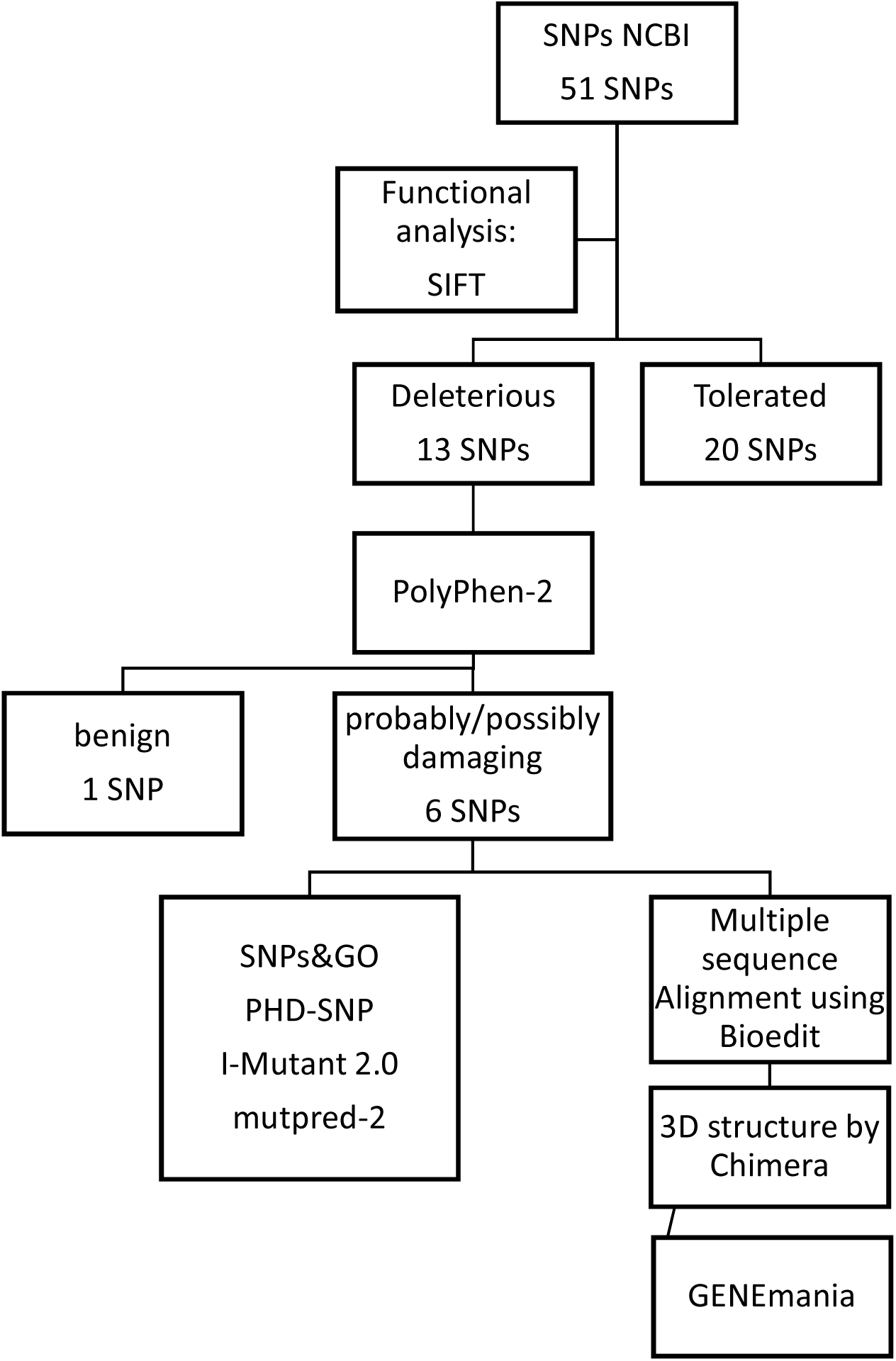
a hierarchy graph showing workflow of *SBDS* SNPs analysis

### Sequence Databases

#### National Center for Biotechnology Information (NCBI)

Is the largest database concerned in providing biomedical and genomic information, it was used in this study to obtain the single nucleotide polymorphisms (SNPs) of the SBDS gene from dbSNP, and reference protein sequences of SBDS family (Homo sapiens, Mus Musculus, Rattus Norvegicus, Felis Catus, Sus Scrofa, Xenopus Laevis) with accession numbers ([NP_057122.2], [NP_075737.1], [NP_001008290.1], [XP_003998651.1], [NP_001231322.1], and [NP_001092159.1], respectively). Adding to that, it was used for obtaining precise information concerning the gene of the study from genebank and nucleotide databases. (Available at: https://www.ncbi.nlm.nih.gov/)

#### ExPaSy

(The Expert Protein Analysis System) is a proteomic database by the Swiss Institute of Bioinformatics (SIB). It provides access to a variety of protein databases and proteomic analytical tools by using protein ID. It has been used to obtain the UniProt entry. ^11,34,34^(Available at: https://www.expasy.org/)

#### UniProt

A world wide database for protein sequences and related information. Data can be collected either by SNPs ID or protein ID which is provided by SIFT server, it provides full results on sequence in the shape of detailed information about protein structure, function, interactions and reported mutations, or simply as FASTA format to be used on other programs (e.g.: SNP&GO, and I-mutant 2.0). (Available at: http://www.uniprot.org/).

### Functional analysis of SNPs

***SIFT*** (Sorting Intolerant from Tolerant) is an online tool to predict whether an amino acid substitution affects protein function based on sequence homology and the physical properties of amino acids. SIFT has been applied to human variant databases and was able to predict mutations (deleterious) involved in disease from (tolerant) polymorphisms which has no ability to alter protein. Thus, cause a disease.The cutoff value in the SIFT program is a tolerance index of ≥0.05. The higher the tolerance index, the less functional impact a particular amino acid substitution is likely to have.^35^ (table 1, 2) (Figure 2) (Available at: http://sift.jcvi.org/)

**Table (1):**
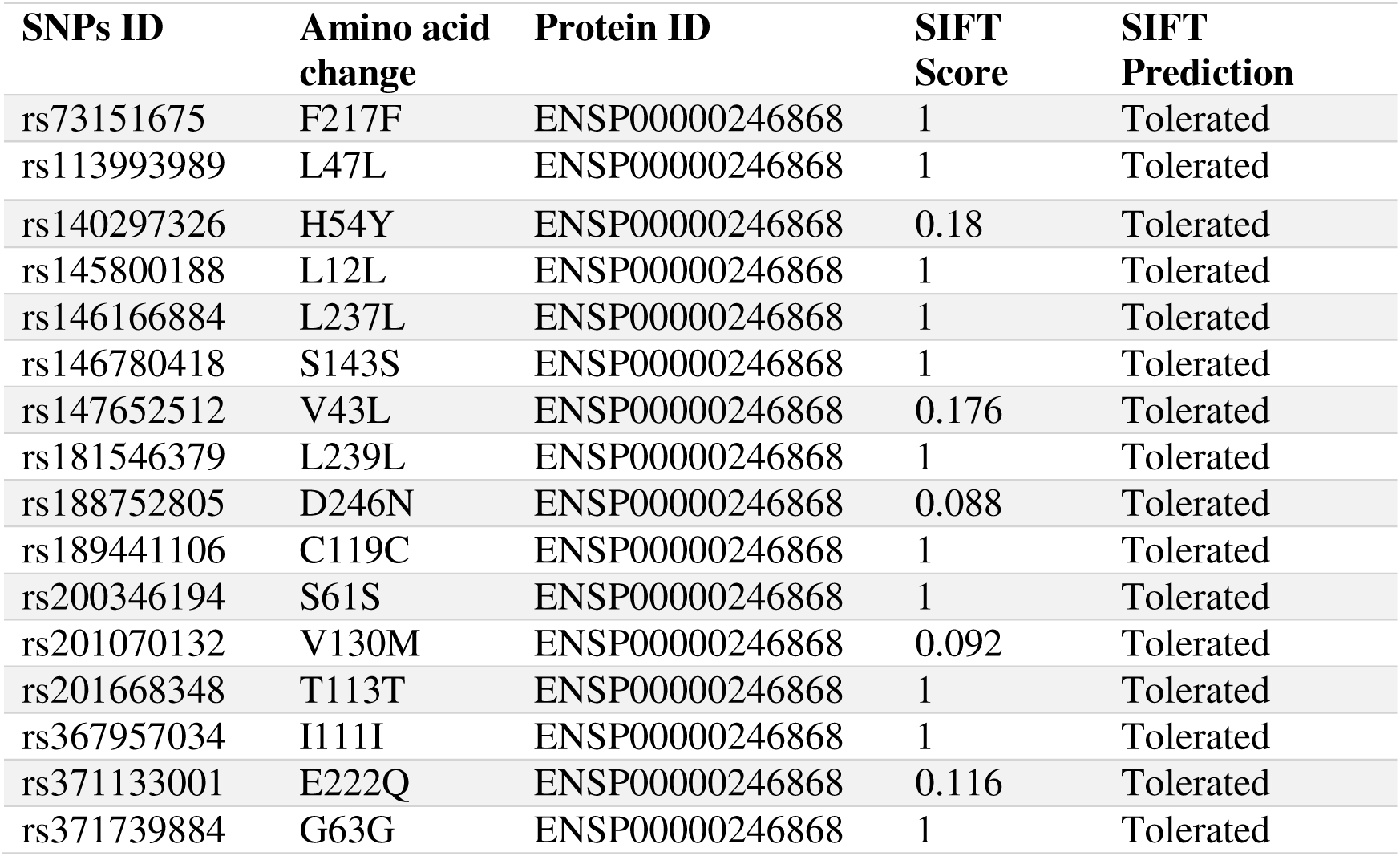
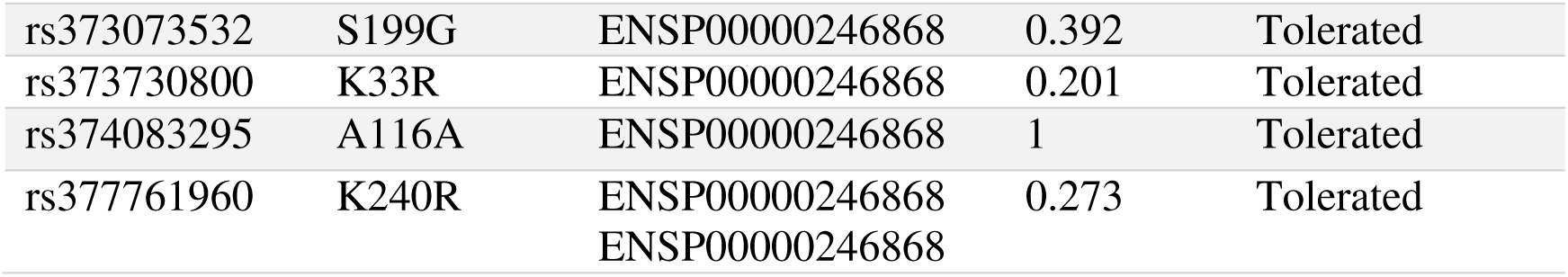
Single nucleotide polymorphisms (SNPs) predicted to be tolerant using SIFT server

**Table (2):** SNPs results using SIFT server and Polyphen-2 server.

| SNP ID | SIFT |  |  | polyPhen-2 |  |
| --- | --- | --- | --- | --- | --- |
|  | Amino acid change | score | Prediction | value | prediction |
| rs11557408 | V17M | 0.002 | Deleterious | 0.999 | Probably damaging |
| rs11557409 | R11H | 0.012 | Deleterious | 0.988 | Probably damaging |
| rs28942099 | N8K | 0.032 | Deleterious | 0.970 | Probably damaging |
| rs79344818 | I212T | 0.017 | Deleterious | 0.145 | Benign |
| rs113993995 | R126T | 0.003 | Deleterious | 0.990 | Probably damaging |
| rs376960114 | F57S | 0 | Deleterious | 0.988 | Probably damaging |
| rs367842164 | A66T | 0.002 | Deleterious | 0.858 | Possibly damaging |

**Figure (2):**
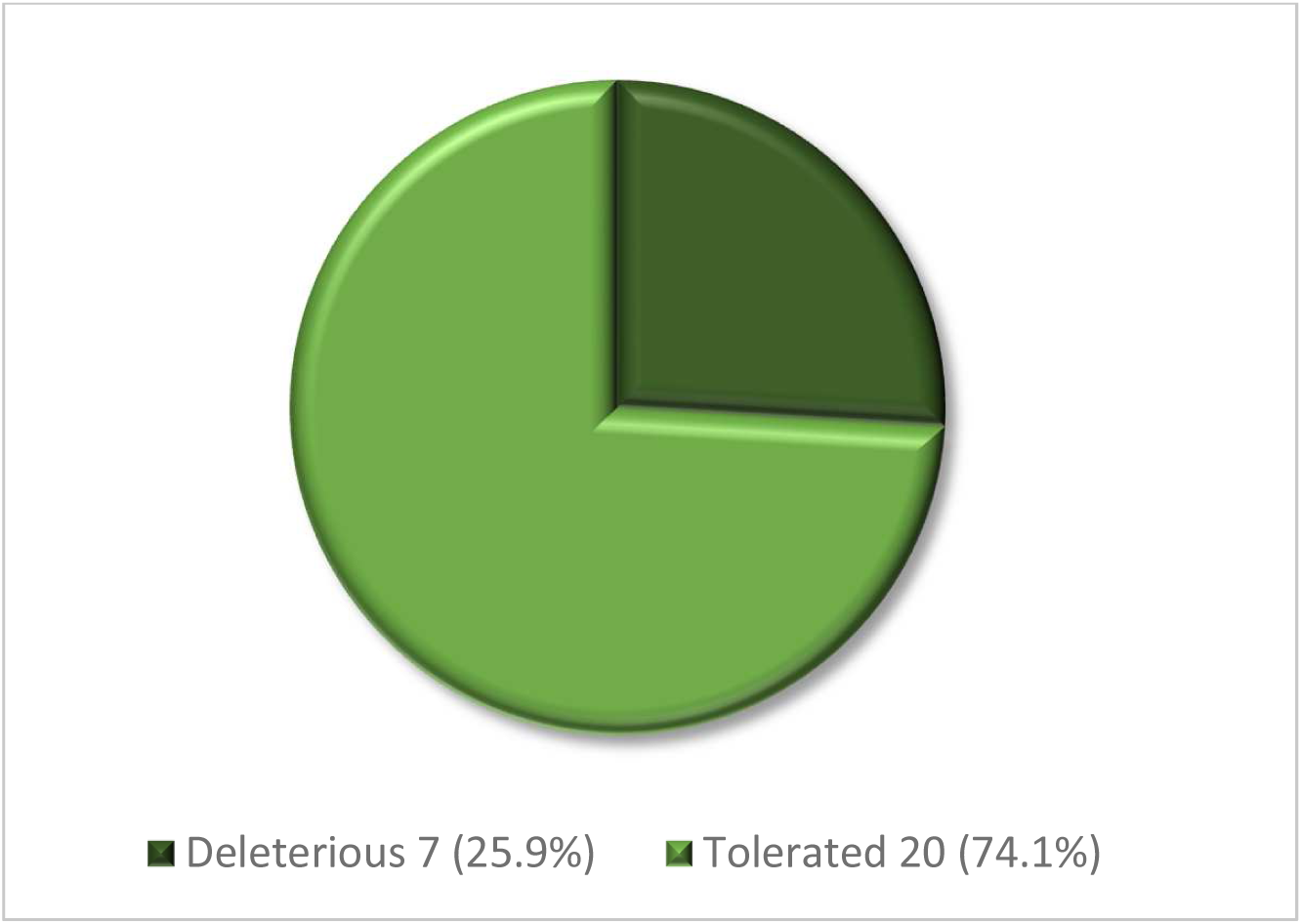
number of SNPs predicted as deleterious and tolerant using SIFT server

#### Polyphen-2

PolyPhen-2 (Polymorphism phenotyping volume 2) is an online translational tool to predict the possible impact of single amino acid substitutions on the function of human proteins using structural and comparative evolutionary considerations, through multiple sequence alignment and protein 3D structure analysis. It also it calculates position-specific independent count scores (PSIC) for each of the two variants, and then calculates the PSIC scores difference between two variants for quantitative assessment of the severity of the effect on protein function. Prediction outcomes could be classified as [benign, possibly damaging or probably damaging] according to the value of PSIC with scores designated as (0.00–0.4), (0.5–0.8), and (0.9 –1) respectively. (Table 2)(Figure 3) (available at: http://genetics.bwh.harvard.edu/pph2/).^7,96^

**Figure (3):**
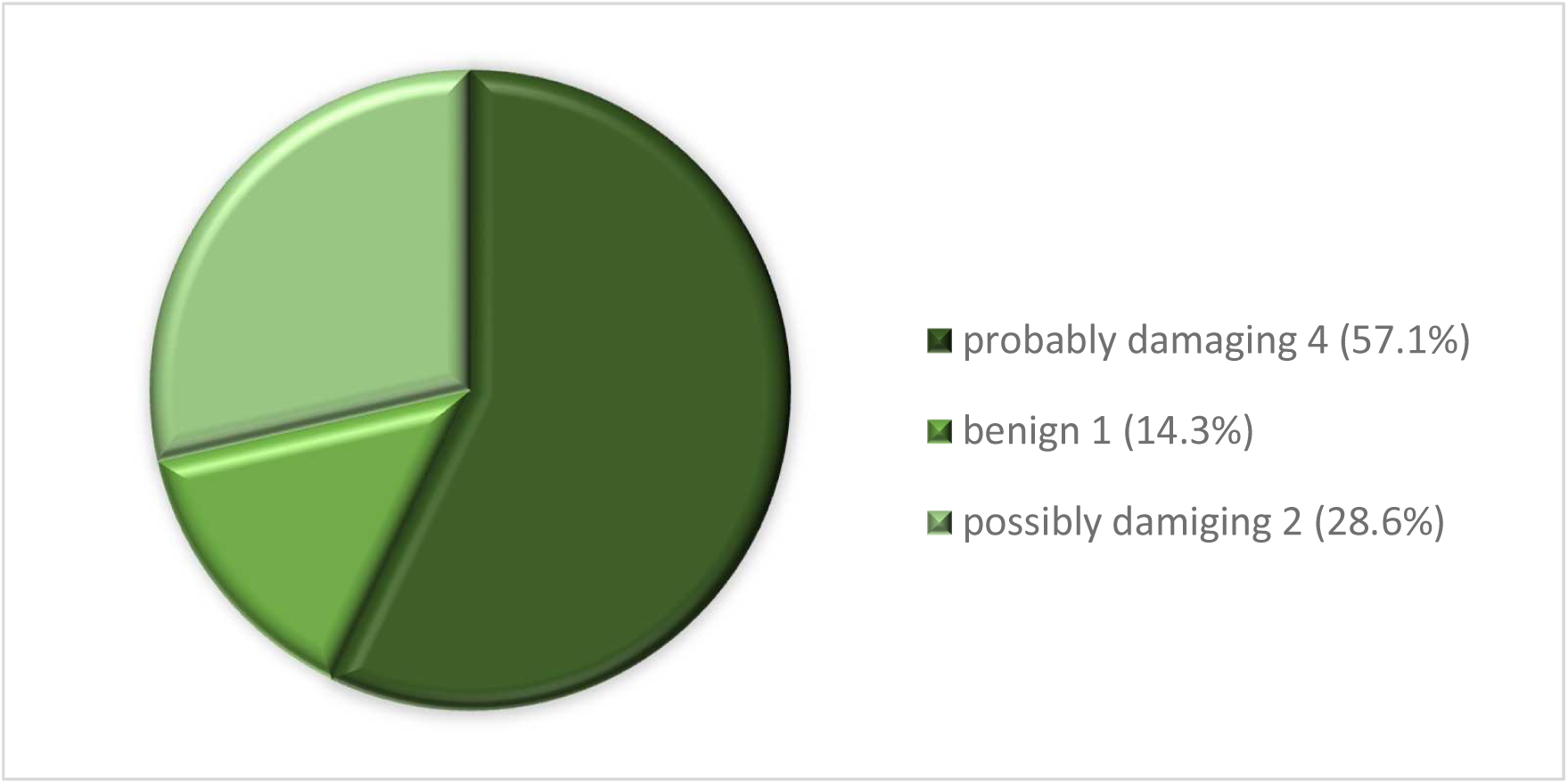
number of SNPs and their disease relationship predictions according PolyPhen-2 server

#### SNPs&Go

It’s an online software or program, It illustrate three results based on three different analytical algorithms, panther result, PHD-SNP result, and SNPs&Go result. All the results are contained in three main parts, the prediction which decides whether the mutation is neutral or disease related, reliability index (RI), and disease probability (if >0.5 mutation is disease causing).^364^(table 3) (figure 4) (Available at: http://snps.biofold.org/snps-and-go/snps-and-go.html)

**Table (3):** Single nucleotide polymorphisms SNPs analysis using SNPs&Go and I-mutant 2.0 server

| SNP ID | Amino acid change | SNPs&GO |  |  | PHD-SNP |  |  | I-mutant 2.0 |  |
| --- | --- | --- | --- | --- | --- | --- | --- | --- | --- |
|  |  | Value | prediction | RI | value | prediction | RI | prediction | RI |
| rs11557408 | V17M | 0.256 | Neutral | 5 | 0.427 | Neutral | 1 | Increased | 2 |
| rs11557409 | R11H | 0.19 | Neutral | 6 | 0.536 | Disease | 1 | Decreased | 8 |
| rs28942099 | N8K | 0.348 | Neutral | 3 | 0.667 | Disease | 3 | Increased | 4 |
| rs113993995 | R126T | 0.291 | Neutral | 4 | 0.514 | Disease | 0 | Increased | 4 |
| rs367842164 | A66T | 0.59 | Disease | 2 | 0.71 | Disease | 4 | Decreased | 7 |
| rs376960114 | F57S | 0.812 | Disease | 6 | 0.871 | Disease | 7 | Decreased | 9 |

**Figure (4):**
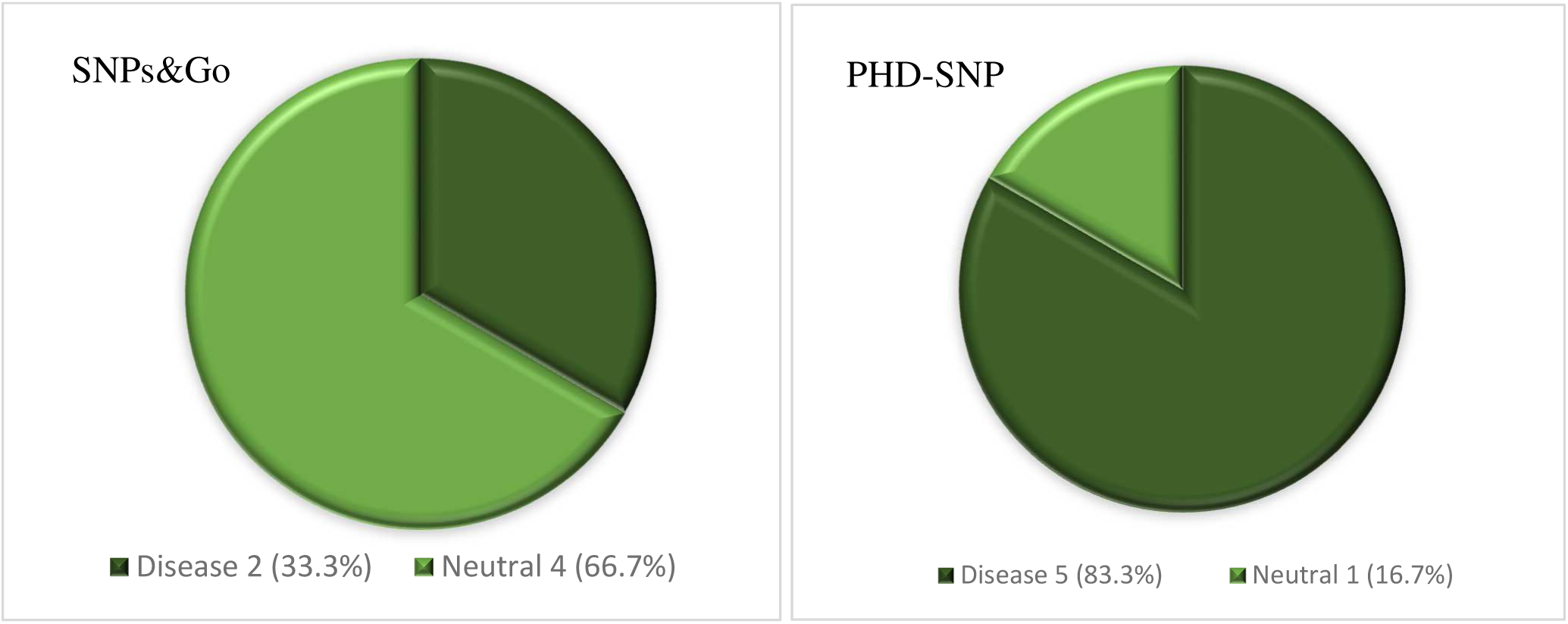
number and prediction of SNPs effect on SBDS protein using SNPs&Go and PHD-SNP softwares

#### PhD-SNP

(Predictor of Human Deleterious Single Nucleotide Polymorphisms). An online support vector machine (SVM) based classifier, uses protein sequence information which can predict if the new phenotype produced after nsSNP can be related to a genetic disease or as a neutral polymorphism.^37^ (table 3) (Figure 4) (Available at: http://snps.biofold.org/phd-snp/phd-snp.html)

#### I-mutant 2.0

Is a support vector machine (SVM) tool for the prediction of protein stability free-energy change (ΔΔG or DDG) on a specific nsSNP. It predicts the free energy changes starting from either the protein structure or the protein sequence. A negative DDG value means that the mutation decreases the stability of the protein, while a positive DDG value indicates an increase in stability. The result show (prediction, reliability index RI, and a DDG value ^38^(table 3) (Figure 5) (Available at: http://folding.biofold.org/cgi-bin/i-mutant2.0.cgi)

**Figure (5):**
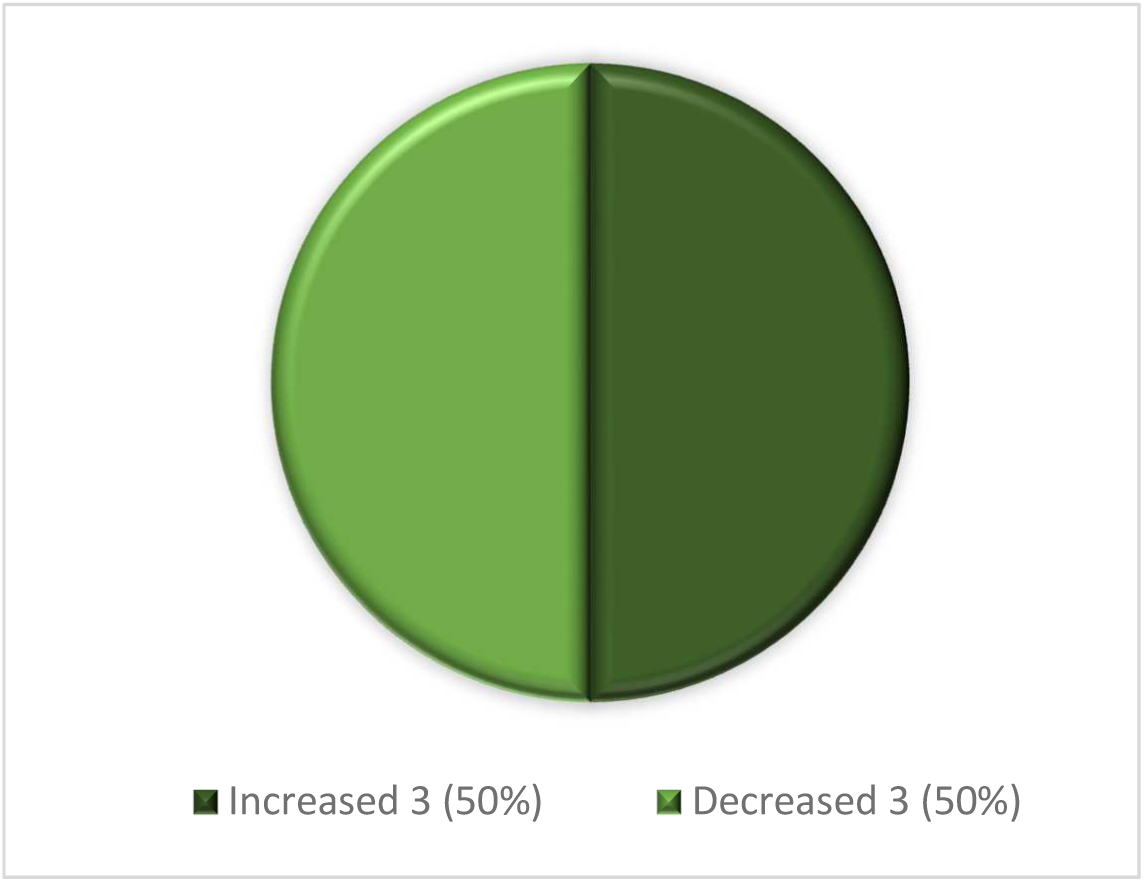
numbers and percentages of SNPs and their effect on protein stability (disease causing mutations alter protein stability either by increasing or decreasing it) using I-Mutant 2.0 server.

### Structural analysis of SNPS

#### MutPred2

Is a tool that detects the effect of missense mutations on protein structure by providing it with the protein sequence in a FASTA format, mutation position, wild and mutant types. The result contains Mutpred2 score which is the probability that the amino acid substitution is pathogenic [>0.50 is considered pathogenic], affected PROSITE and ELM motifs, and a table with the structural changes occurred whether loss/gain of a helix, strand or loop. Also, gives other structural changes information if any and a P-value.^39,40^ (table 4) (Available at: http://mutpred2.mutdb.org/index.html)

**Table (4):** Showing structural effects of mutation based on Mutpred2 tool.

| SNPs ID | UnitProt ID | Amino acid change | Mutpred2 Score | PROSITE/ELM motifs |
| --- | --- | --- | --- | --- |
| rs11557408 | Q9Y3A5 | V17M | 0.645 | — |
| rs11557409 | Q9Y3A5 | R11H | 0.346 | — |
| rs28942099 | Q9Y3A5 | N8K | 0.857 | ELME000052<br>ELME000063<br>ELME000153<br>ELME000159<br>ELME000233<br>ELME000358 |
| rs113993995 | Q9Y3A5 | R126T | 0.644 | ELME000100<br>ELME000108<br>ELME000136<br>ELME000142<br>ELME000159<br>ELME000233<br>PS00005 |
| rs367842164 | Q9Y3A5 | A66T | 0.753 | ELME000002<br>ELME000064<br>ELME000220<br>PS00005<br>PS00006 |
| rs376960114 | Q9Y3A5 | F57S | 0.912 | ELME000052<br>ELME000053<br>PS01267 |

#### Chimera 1.13.1

Is a software program for analysis of molecular structures, visualizes protein in 3D structure of the protein and then change it between native and mutant amino acids with the candidate to display the impact that can be produced. Chimera accepts the input in the form of PDB ID or PDB file. ^41–43^ (Figure 6) (Available at: http://www.cgl.ucsf.edu/chimera)

**Figure (6):**
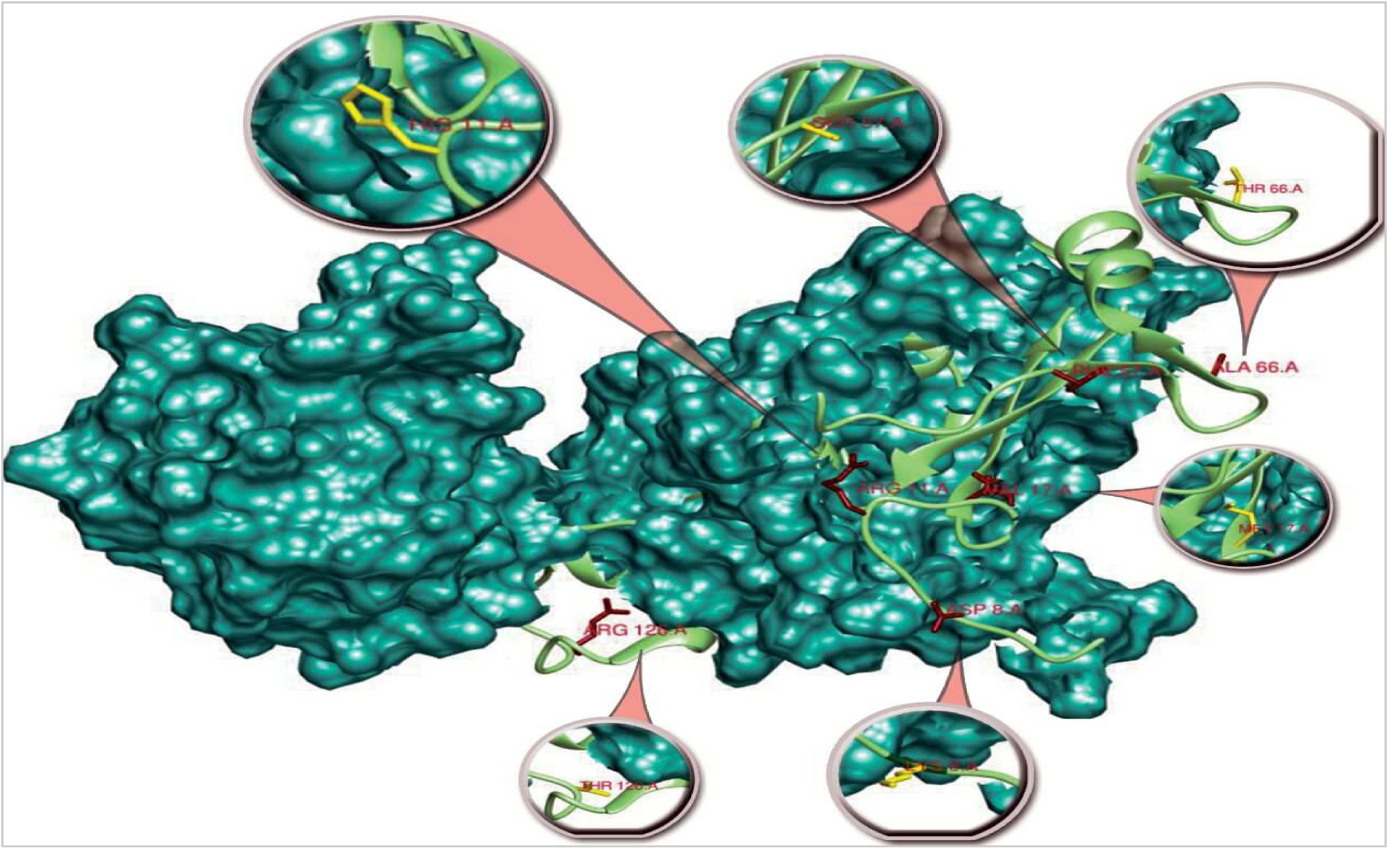
3D structure of SBDS protein showing the six disease causing mutations, wild types (Red) and mutant types (Yellow) using Chimera 1.13.1.

### Multiple Sequence Alignment

Using **Clustal W** package, **Bioedit** software which is a nucleic acid/protein alignment and BLAST tool. It was used to align six of SBDS protein family to illustrate areas of conservation and identities^44^. Areas of high conservation are the areas associated with protein functions. Thus, mutations located at highly conservative regions (motifs) are more capable of altering protein structure and function and causing pathology than mutations at areas of low conservation.^45–47^ (Figure 7)

**Figure (7):**
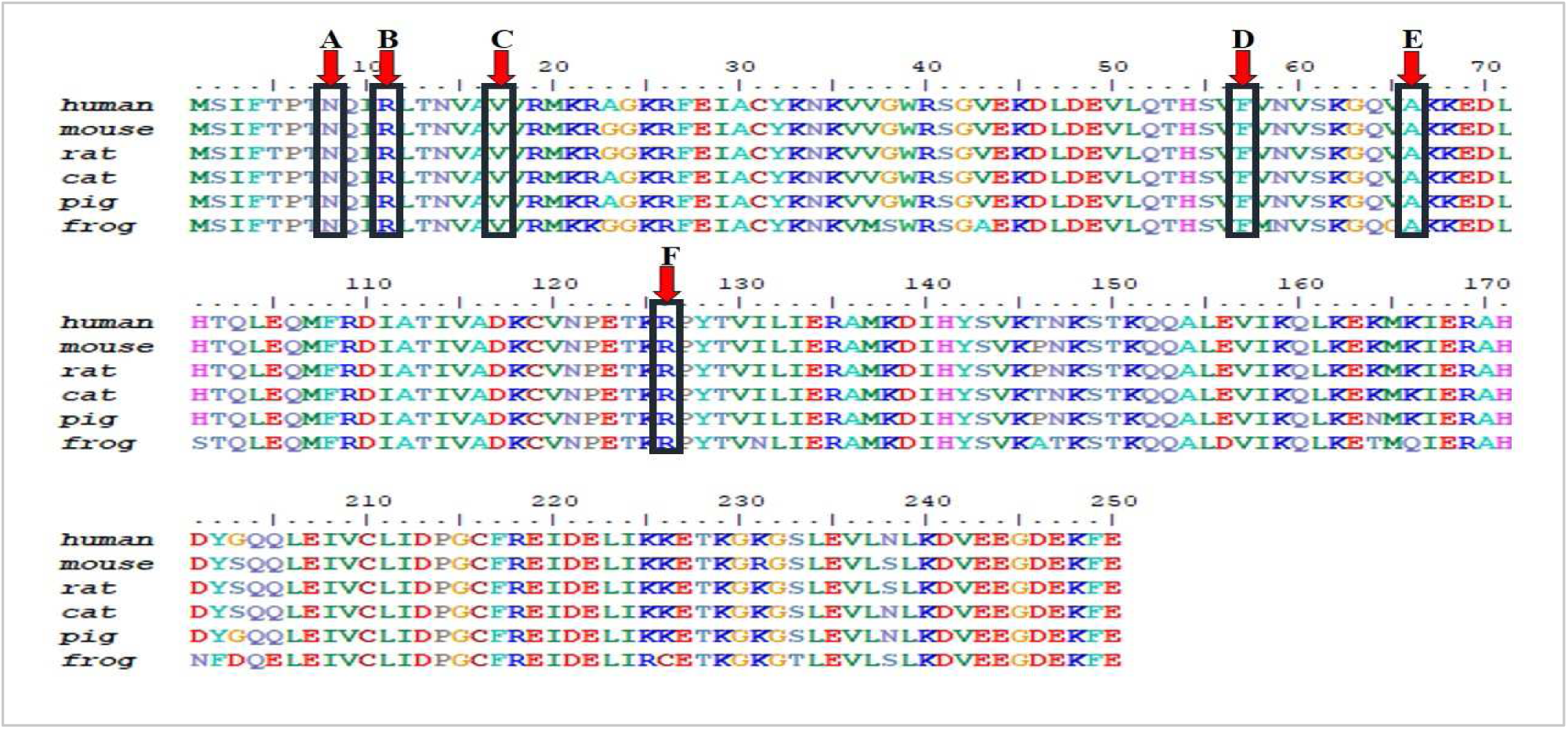
Multiple sequence alignment of the SBDS Protein family, shows mutations (**A.** rs28942099 (N8K), **B.** rs11557409 (R11H), **C.** rs11557408 (V17M), **D.** rs376960114 (F57S), **E.** rs367842164 (A66T), **F.** rs113993995 (R126T)) are located at areas of high conservation using Bioedit software.

### Prediction of gene functions and interactions

Using **GeneMANIA version 3.6.0** which is an online server that finds genes which are related to an input gene, through functional association data. These data provide gene to gene (co-expression), gene to protein (physical interactions), pathways the gene is a part of and co-localization of the gene with a high accuracy.^48^ The genes co-expressed and/or interacting with SBDS gene and their descriptions are shown in (table 5) (figures 8, 9). (Available at: http://www.genemania.org)

**Table (5):**
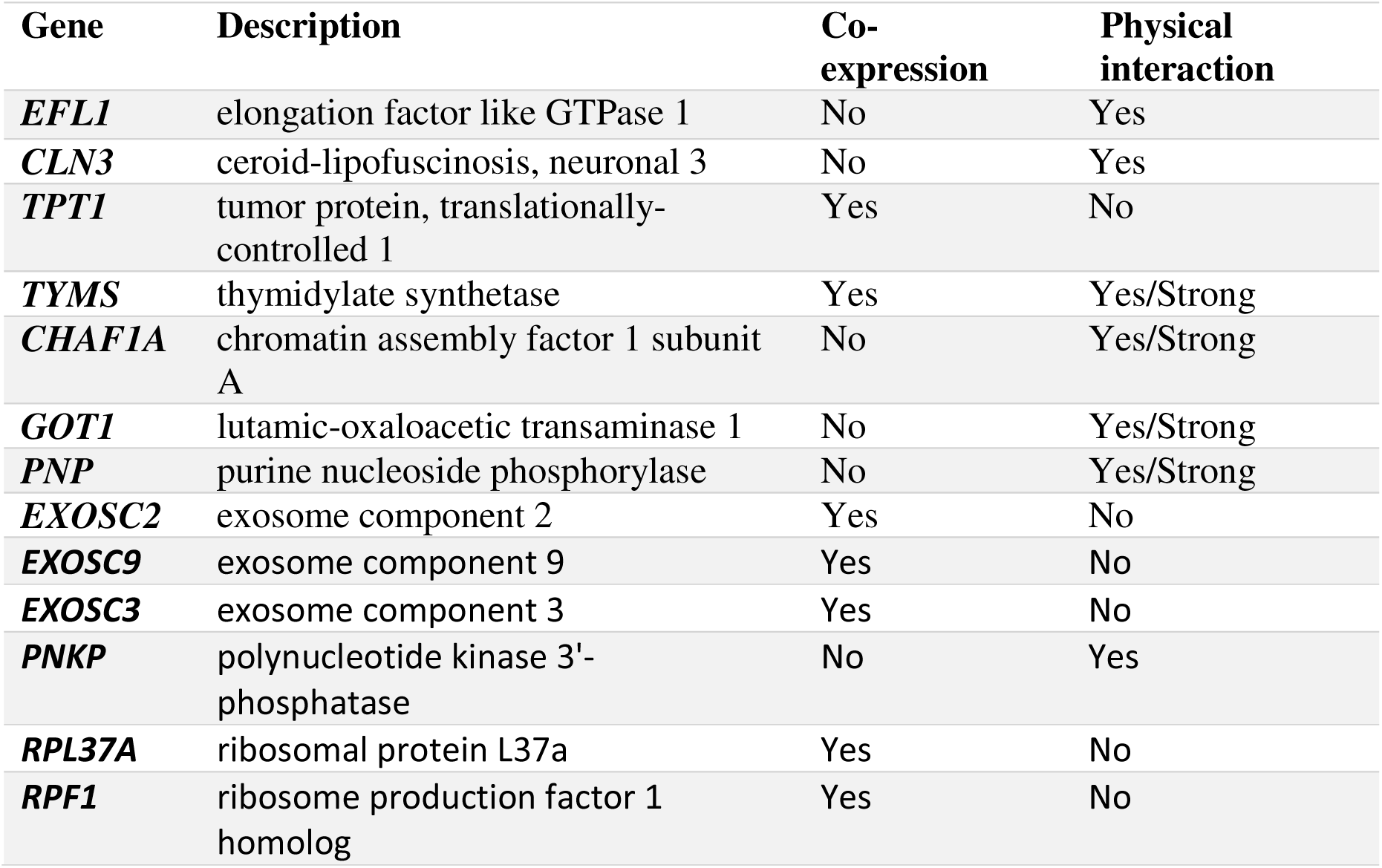
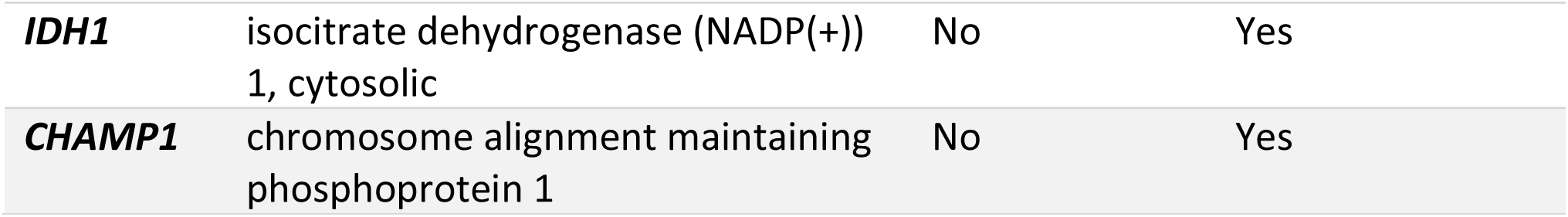
list of genes co-expressed/physically interacting with *SBDS* gene and their description:

**Figure (8):**
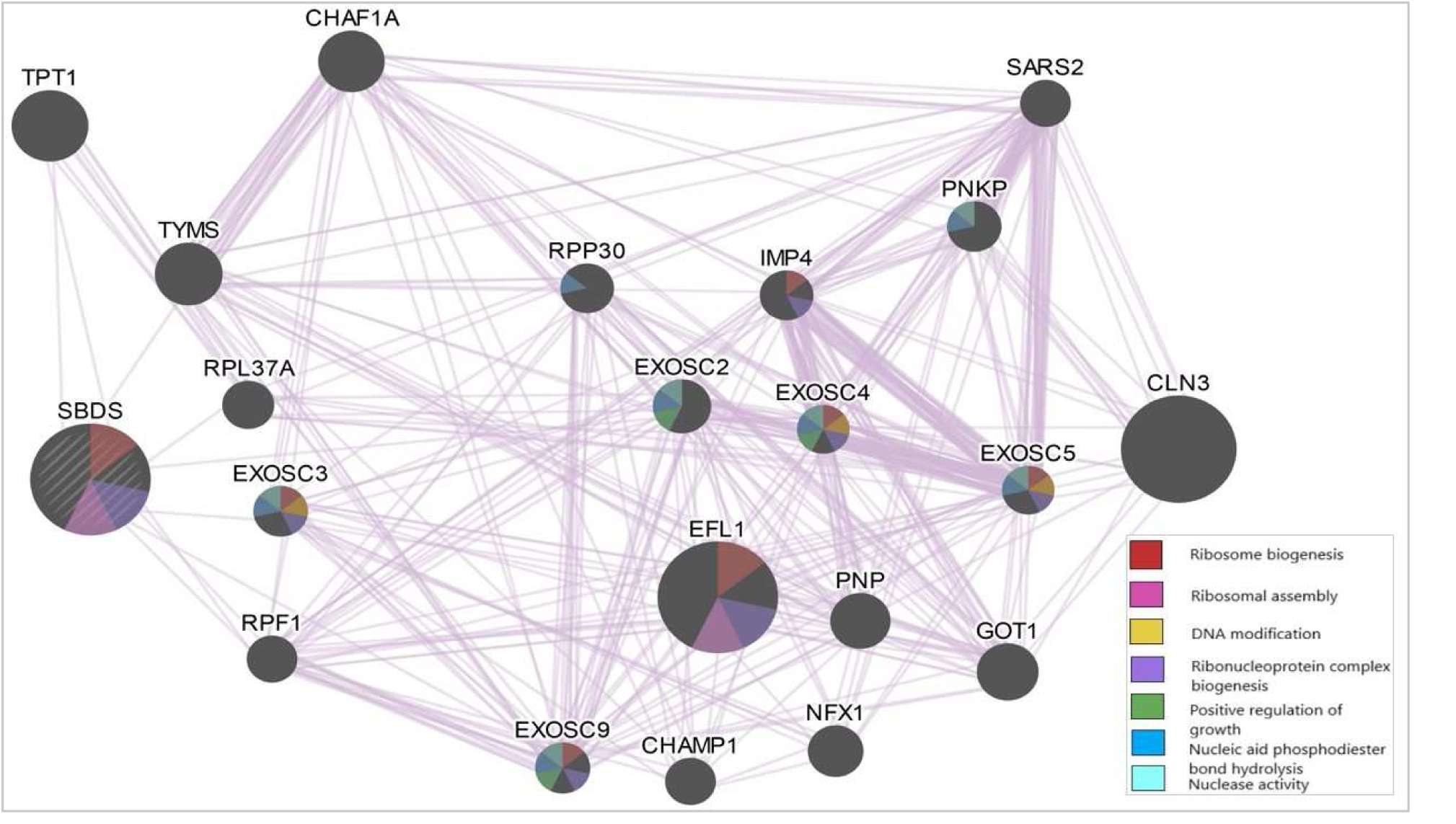
GENEmania. network showing genes co-expression patterns with *SBDS* gene, color codes indicates functions shared by genes.

**Figure (9):**
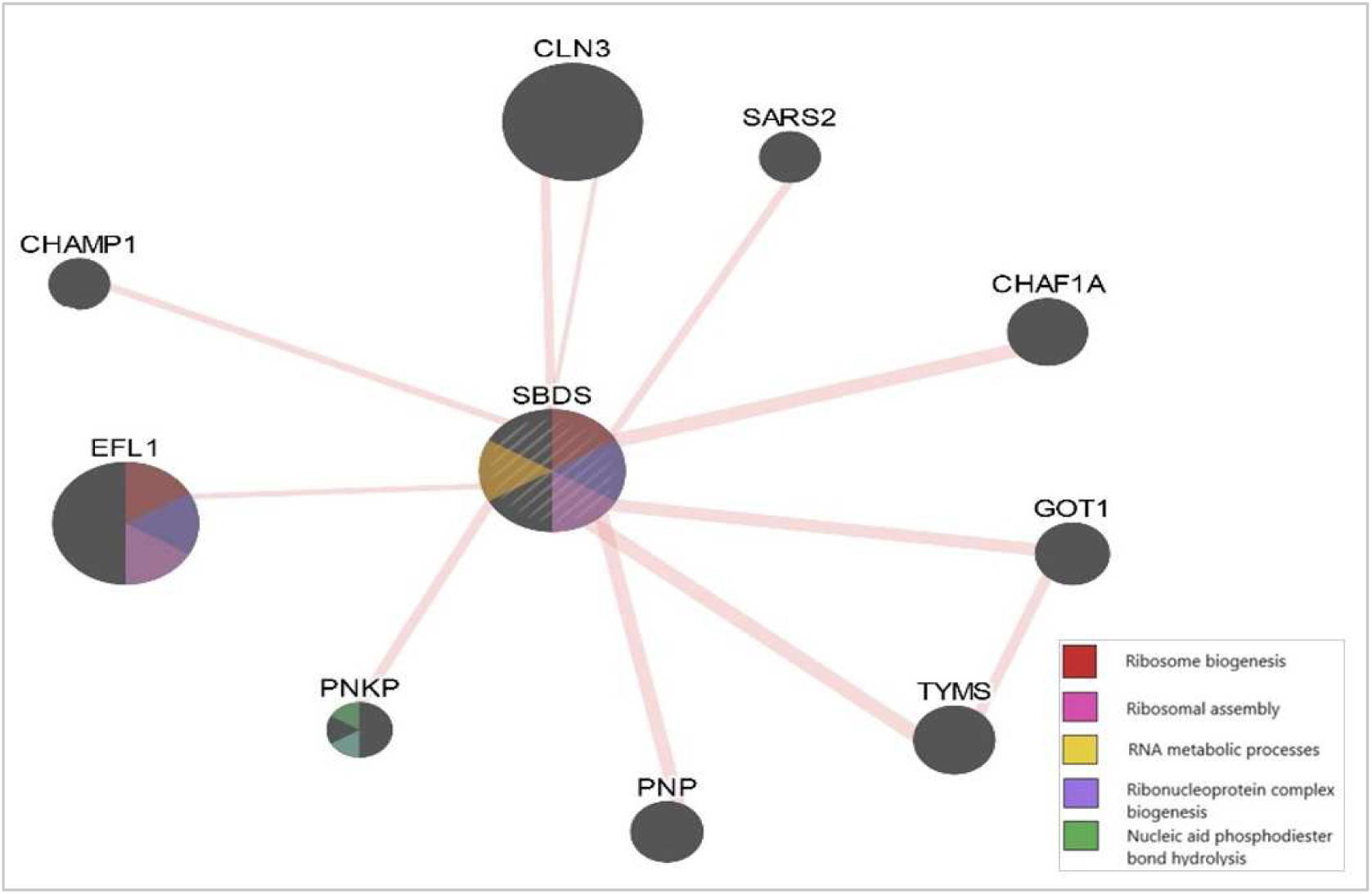
GENEmania. network showing genes physical interaction patterns with *SBDS* gene, color highlights indicates shared functions.

## Results

## Discussion

Six SNPs were found to be the most disease causing SNPs (rs11557408, rs11557409, rs28942099, rs113993995, rs367842164, rs376960114) ((V17M), (R11H), (N8K), (R126T), (A66T), (F57S)) indicating they have the ability to change the SBDS protein structure and functions; thus causing SDS.

The following SNPs with IDs (rs11557408, rs11557409, rs367842164, and rs376960114) and mutations of ((V17M), (R11H), (A66T), and (F57S), respectively) have been reported to be deleterious by SIFT server, and probably/ possibly damaging according to PolyPhen-2. Mutations of both (A66T) and (F57S) were predicted to be double disease causing by SNPs&Go and PHD-SNP and showed decreased protein stability, while (R11H) was disease causing by PHD-SNP, neutral by SNPs&Go and showed decreased protein stability, and (V17M) was neutral by both with elevated protein stability. However, the four SNPs were located on highly conservative region of the SBDS protein as previously shown in the multiple sequence alignment with SBDS protein family and were not reported by any previous study regarding this gene.

Mutations of Asparagine into Lysine at position 8 (N8K) (SNP ID rs28942099), Arginine into Threonine at position 126 (R126T) (SNP ID rs113993995), Isoleucine into Threonine at position 212 (I212T) (SNP ID rs79344818) were found by Boocock’s et el 2002 study, these findings are in agreement with our findings, the mutations were predicted to be disease causing according to SIFT. (rs28942099, and 113993995, (N8K) and (R126T), respectively) were probably damaging by PolyPhen-2 servers, neutral according to SNPs&Go and disease causing according to PHD-SNP with an increased protein stability according to I-mutant 2.0, while (rs79344818) (I212T) was benign and had not further analyzed. Additionally, according to Mutpred2 prediction mutation of SNP (rs28942099) (N8k) led to structural changes including a gain of strand. Mutation of (R126T) (SNP ID rs113993995) was also reported by another study Finch et al. 2005 to be defective using protein expression and purification techniques which comes also in agreement to this study’s findings.^17,31^

Mutation at position 33 of the wild amino acid Lysine was reported in a study by Carvalho et al. 2014, using Array comparative genomic hybridization and Southern blotting detected mutation in a SDS patient, and study by Shammas et el.2005 using western blotting and flowcytometry techniques.^32,30^ However, different mutant types occurred each time (Threonine (K33T), Glutamic acid (K33E), respectively). A similar mutation in the same position 33 and wild type Lysine converted into Arginine (K33R) was found in study by Boocock et el 2002 using southern hybridization and RT-PCR techniques in SDS affected individuals.^17^ This mutation is similar to the one obtained from NCBI database in this study (SNP ID rs373730800) (K33R) and had been predicted to be tolerated by SIFT server and had not been analyzed furthermore, which contradicts the previous studies results.

As previously mentioned in the introduction, *SBDS* gene plays an important role in ribosomal processes, most importantly, ribosomal biogenesis through (physical interaction with EFL1, and co-expression with EXOSC3, and EXOSC9), positive regulation of cell growth (co-expression with EXOSC2, EXOSC4, and EXOSC9), ribosomal assembly (through interaction with EFL1 only), and undergoing apoptotic and cellular clearance processes through (co-expression with TPT1, Exosome components Two, three, nine and physical interaction with GOT1).^49–52^ According to GENEmania report, *SBDS* had strong physical interaction with nine genes, mainly (TYMS, CHAF1A, GOT1, and PNP). A lesser degree of physical interaction with both (RPP30, and CHAMP1). In addition, it had been co-expressed with eight genes (TPT1, TYMS, EXOSC2, EXOSC3, EXOSC5, EXOSC9, RPL37A, and RPF1).

This study encountered possible new functions of the *SBDS* gene, which is predicted to have new DNA related functions through GENEmania, as it has been both co-expressed and strongly interacting physically with (TYMS – a gene which plays an important role in pyrimidine metabolism and synthesis of dTTP and has polymorphisms associated with developing neoplasia-, CHAF1A –the chromatin assembly unit in DNA replication an repair-, and PKNP –responsible of DNA repair-)^53–55^. It was also found that SBDS gene is co-expressed with RBF1 gene (a gene mainly expressed in the bone marrow which is responsible of ribosomal assembly, and RNA processes)^56^. This suggests that, mutations in the SBDS gene could result in impaired expression of RBF1 gene resulting in patients developing variable levels of cytopenias. In addition to that, SBDS gene has a co-expression with EXOSC3 gene (which is partially responsible of muscle movement), impairment in expression of SBDS gene would affect the proper expression of EXOSC3 as well. Thus, this could also be used as an explanation of SDS patients developing skeletal abnormalities and failure to thrive.^57^

The limitation of this study was using computational analysis only, we recommend wet lab analysis of the four novel mutations reported in this study, and clinical and laboratory analysis of the new predicted gene functions.

The most important findings in this study were four mutations predicted to be disease causing which have not been reported previously, (rs11557408 (V17M), rs11557409 (R11H), rs367842164 (A66T), and rs376960114 (F57S)) in addition, predicted functions of SBDS gene which are DNA processes related, RBF1 affected expression leading to cytopenias and EXOSC3 affected expression leading to failure to thrive.

## Conclusion

This study uses insilico tools which are important diagnostic tools, they are necessary to guide molecular geneticists, they balance between the time consumption, low cost required and are more consistent. Not only that, but insilico technology proved its efficiency in the field of therapeutics as well. For conventional drug, vaccine designing and cell therapy methods consumes prolonged time, man power, cost and raises toxicity and safety issues, however, computational tools have been developed to design immunotherapy as well as peptide-based drugs and efficient regenerative medicine tools and proved to be a catalyst in drug and vaccine design with no safety issues at all.

